# SVarp: pangenome-based structural variant discovery

**DOI:** 10.1101/2024.02.18.580171

**Authors:** Arda Söylev, Jana Ebler, Samarendra Pani, Tobias Rausch, Jan O. Korbel, Tobias Marschall

## Abstract

The linear human reference genome that we use today does not represent the haplotypic diversity of the global human population. This raises bias in genomic read alignment and limits our ability to call large structural variations (SV), especially at highly polymorphic loci. Thus, many SV alleles remain unresolved. Recent efforts to transition to a graph-based reference genome resulted in the generation of the first draft human pangenome reference, but tools to call SVs relative to the pangenome reference are presently lacking. In this study, we present the SVarp algorithm, aiming to discover haplotype resolved SVs on top of a pangenome reference using long sequencing reads. SVarp outputs local assemblies of SV alleles, termed svtigs, instead of a VCF file of SV breakpoints, which we propose as a general exchange format allowing for flexible downstream analyses. In order to assess the accuracy of svtigs, we used simulated and real human genomes. Simulations allowed us to make exact breakpoint comparisons against the true callsets. We observed ∼96% recall with deletions, insertions and duplications larger than 1,000bp, showing that SVarp can reliably detect genomic structural variants not yet represented in the graph. On the other hand, we compared SVarp output for ONT sequencing data at 20X coverage against independent genome assemblies of the same samples and found that ∼82% of our svtig predictions are validated by the assemblies by a match with more than 85% sequence identity. SVarp was implemented using C++ and its source code is available at https://github.com/asylvz/SVarp under MIT license.

## 1. Introduction

Genomic analysis has mainly been based on using a linear reference genome since the release of its initial draft in 2001 [1]. However, this approach is inadequate to represent the genetic diversity of a species [2–5]. Given the fact that the human reference genome (e.g., GRCh38) harbors only a single representative scaffold for each chromosome as the primary sequence, using it as a reference genome comes with severe limitations, including population-specific read mapping biases.

While most studies have focussed on single nucleotide polymorphisms (SNPs) and small insertions and deletions (Indels) in the past, there is an increasing interest also in structural variants (SVs), which are an important contributor to human phenotypes in general and to genetic diseases in particular [6–12]. They are known to affect more nucleotides and have higher impact on gene functions compared to SNPs or Indels [13]. We have witnessed two significant advances in genomics recently: (1) developments in sequencing technologies that enabled the cost-efficient generation of much longer reads (>20 kb) with an error rate close to short Illumina reads [14–17]; (2) gaps in the human reference genome have been closed with the completion of the first telomere-to-telomere assembly (T2T-CHM13) [18]. Despite these advances, the problem of SV discovery has still not been fully resolved, especially in low mappability and repeat-rich regions [19–21]. While SV discovery using long reads increased the accuracy of the algorithms in characterizing variants due to increased mappability of repeats compared to short reads, there are still problems adherent to the reference bias resulting from using a single linear reference genome. One of the main reasons is that the algorithms (either short-read [22–25], long-read [26–28] or assembly based [6,29]) rely on mapping reads/contigs to a reference assembly and lack of alternative alleles in the linear reference impacts read alignment accuracy, thus limiting the number of detectable SVs.

With the recent studies, there is an ongoing endeavor to combine multiple haplotypes in a graph structure—a transition to pangenomes—, aiming to address the reference bias [3,20]. With the efforts of the Human Pangenome Reference Consortium (HPRC), the first draft human pangenome reference has been generated using 44 phased diploid assemblies, corresponding to 88 haplotypes [4]. In parallel with this effort, pangenomic methods have started to emerge [3,30–33]. These studies showed that there is a significant improvement in SV genotyping using short-reads with pangenomes compared to a linear reference, therefore the pangenome encodes known SVs and there has been great progress on re-detecting these known SVs [34,35]. Despite this progress, there is still a lack of tools able to discover novel SVs from read alignments to a pangenomic reference, that is, to find SVs that are not yet represented in the pangenome graph.

Here we present SVarp (***S****tructural* ***Var****iations in* ***P****angenome*s) to address this gap by calling SVs on graph genomes using third generation long sequencing reads (either ONT or HiFi). This enables us to find additional SVs that are currently missing, including SVs on top of alternative sequences present in the pangenome but not in a linear reference. With SVarp, our aim is to call novel phased variant sequences, which we call svtigs, rather than a variant call set. This way, the variant representation is not tied to a single linear reference and allows for flexible downstream workflows that derive variant calls from these assemblies, either relative to a pangenome reference or, if desired, relative to a linear reference. The svtigs can furthermore serve as a basis to amend a pangenome graph.

## 2. SVarp algorithm

SVarp processes sequencing reads aligned to pangenome graphs (GAF format) in five steps. These are depicted in the workflow (Figure 1) and briefly described below.

**Figure 1.**
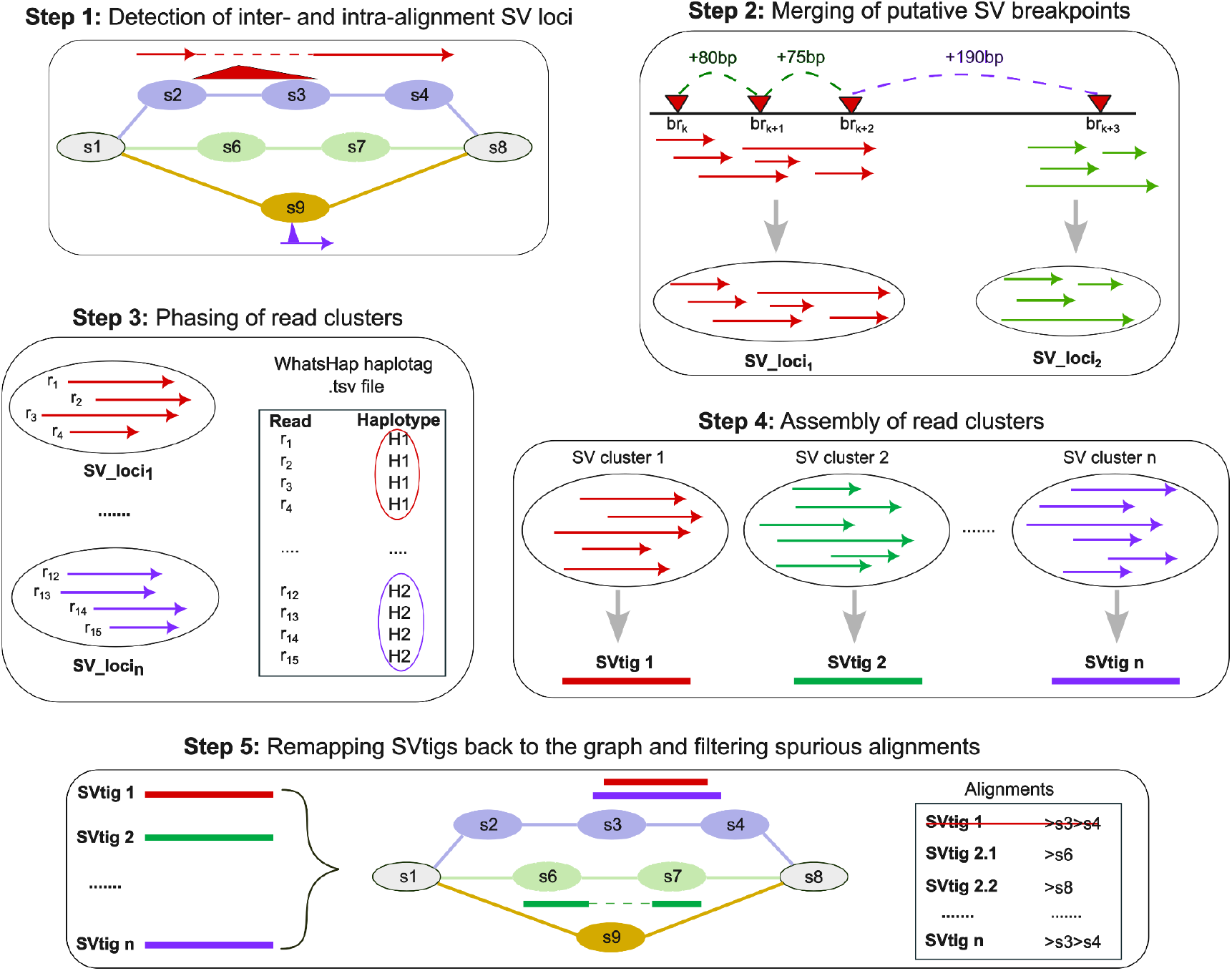
Overview of the SVarp algorithm, which uses a pangenome graph (top left with nodes s1 to s9) and alignments to it (horizontal arrows) as input. Large insertions/deletions in the alignment indicate putative SV breakpoints and are shown as red triangles. Breakpoints, labeled as “br”, below a distance-threshold are merged (Step 2, green dashed line). After phasing (Step 3), clusters are assembled to generate svtigs (Step 4) and spurious svtigs are filtered after remapping them back to the pangenome (Step 5).

### 2.1 Formation of read clusters

In the first step, given the alignments to a pangenome graph, we find putative variant sites inter- and intra-alignment as follows:

1. Intra-alignment variations are defined as >50 bp insertions (“I”) or deletions (“D”) parsed from the CIGAR string of the GAF file. It is straightforward to detect these variants because the SV is contained within the read as shown in Figure 2 Case 1. Of note, the sensitivity to detect intra-alignment SVs, is influenced by length and error-rate of the reads combined with the alignment method and its parameters. We therefore additionally detect inter-alignment signals as described below.
2. Inter-alignment signatures for SVs arise when a SV is large compared to the read length or represents a complex rearrangement. In such cases, reads overlapping the variant can be clipped into multiple partial alignments. Thus the variant is not encoded in the CIGAR. Here, we consider the end of the initial alignment and the beginning of the other alignments as putative breakpoint loci. If there is a deletion within the read, a region on the contig that the read is supposed to map, is left unmapped (Figure 2 Case 2, under Deletion). On the other hand, for insertions, the inserted segment within the read is left unmapped and the other segments are mapped adjacent to each other (Figure 2 Case 2, under Insertion). There is also a third case for insertions where the read harbors a very large inserted segment such that it is left unmapped (Figure 2 Case 3, under Insertion). In order to detect these types of variations, we cluster those reads separately.

**Figure 2.**
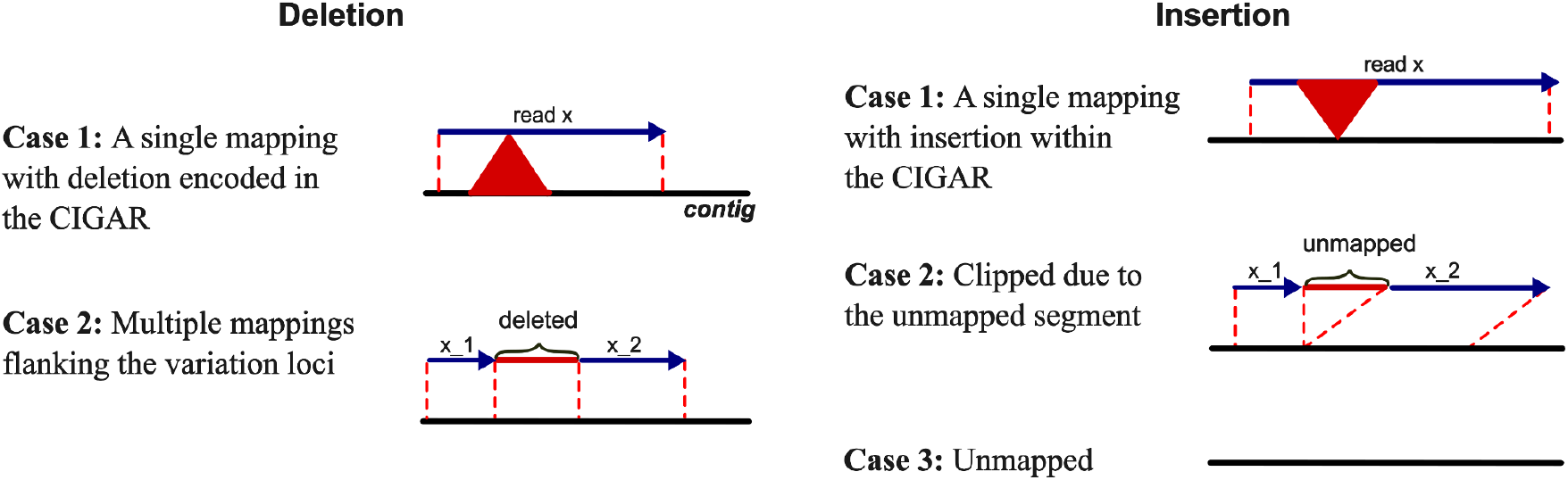
When a “read x” is aligned to a contig, there are 2 cases if the read spans a deletion (left) and 3 cases if it spans an insertion (right). A read can map as a single alignment (Case 1) or split into multiple alignments as “x_1” and “x_2” (Case 2). On the other hand, if “read x” harbors a large insertion, then it may be left as unmapped by the aligner (Case 3).

Our approach involves clustering the reads that contribute to each breakpoint—allowing a read to be present in more than one cluster—assuming each cluster as an SV candidate. In order to avoid alignment artifacts, we merge any two breakpoints that are in close proximity. They are supposed to be pointing to the same variant (Figure 1, Step 2). We note that it is always possible that two very close variants might both be real, however, these will not be missed by merging because the variation assemblies that we generate in the end will harbor both variants. By default, we use 100 as the distance threshold, however for lower coverage genomes, we observed that higher values (e.g., 250) increase recall (see Results Section for comparison).

### 2.2 Phasing

Having the final SV breakpoint set and the set of reads contributing to the SV sites, we integrate read-level haplotype information and divide the read clusters based on the haplotype as depicted in Figure 1, Step 3. This is done after we filter read clusters that have very low read support (default=5).

In order to generate phased variants with SVarp, optional phasing information needs to be given as input (using “--phase file_name”). The expected format is the one generated by WhatsHap haplotag (.tsv file) [36]: “<read_name, haplotype, phase_set, chromosome>“. The approach involves scanning the reads contributing to each cluster and dividing each SV into three phase sets as “Haplotype 1”, “Haplotype 2” and “Untagged”. Thus, reads contributing to each SV are divided into these sets before the assembly step and are assembled separately.

### 2.3 Assembling the final clusters

As shown in Figure 1, Step 4, we assemble each read cluster to generate consensus sequences, which we call an “svtig”. Here, we use haplotagged read clusters as input to the assembly, i.e., reads divided into three sets as input described in Section 2.2. For each such cluster, we apply read-depth based filtering before the assembly, in order to filter spurious SV signals. To this end, we compute the average read depth across all contigs in the pangenome graph, which we call expected read depth λ, as well as the observed read depth x of each cluster. We assume that very high and very low observed read depth compared to the expected is an indication of a complex region where either the pangenome graph representation is wrong or incomplete, or where the alignments of ONT reads are ambiguous due to long repeats. Thus, we filter SV haplotypes with number of reads > 2 * λ as a high coverage region and with < 1/5 * λ as a low coverage region. Then we assemble SV haplotypes that have enough depth by using the wtdbg2 assembly algorithm [37]. Finally, we concatenate all the assemblies into one fasta file to be used as input to the realignment and filtering step.

### 2.4 Realignment and filtering

In the final step (Figure 1, Step 5) of our workflow, we filter spurious and duplicate svtigs by mapping them back to the genuine pangenome graph using GraphAligner [32]. According to our observations, GraphAligner often generates more contiguous alignments compared to Minigraph [30], that is, a read aligned using Minigraph is clipped into multiple alignments (possibly due to its seeding strategy), whereas the same read is mapped in one piece with high sequence identity with GraphAligner. One such scenario is shown in Supplementary Figure S1. While GraphAligner is slower than Minigraph (Supplementary Table S1), the number of unfiltered svtigs that we input to the realignment step is low compared to a classic genome alignment process and we observed that the performance of SVarp is not affected much in terms of runtime (See Section 3.3 for runtime and memory consumption analysis).

Although GraphAligner is robust in finding the correct alignment loci, it performs edit-distance based alignments, leading to biologically implausible placement of gaps. In order to overcome this, we use the wavefront alignment algorithm (WFA) [38,39] and perform an affine-gap cost based realignment of each svtig to the path in the graph output by GraphAligner (see Supplementary Figure S2). Ultimately, we remove low mappability and secondary alignment svtigs as well the ones that have a single alignment but the SV is not found in the cigar string as “I”, “D” or “X”. For duplicate removal, we check the svtigs that have >90% overlap ratio and keep the larger one.

## 3. Results

As a first assessment of SVarp, we used a haploid simulated genome with small and large size SVs. Additionally, we tested our algorithm using ten genomes from the 1000 Genomes Project (1KG) cohort sequenced on the ONT platform with various depths of coverage.

### 3.1 Accuracy of SVarp using simulated genomes

For the simulation experiments, we used VISOR [40] to simulate reads with an average length of 10 kbp that harbor small and large SVs randomly generated from a uniform distribution in the range [50, 1000] and [1000, 10000] bps respectively. This produced FASTQ files for both small and large variant types of insertions, deletions and duplications that harbor 276 implanted SVs each. Then we aligned these six sequence files to the HPRCv1.0 T2T-CHM13 Minigraph pangenome graph [4,41] separately using the Minigraph sequence-to-graph aligner [30] with default parameters (except “--vc” which outputs alignment positions based on unstable coordinate system) to generate GAF alignment files for each different case. Finally, we ran SVarp using these GAF files as input and generated SV assemblies (svtigs). In order to evaluate the performance of our algorithm, we called specific variation breakpoints with respect to the linear T2T reference genome using an assembly based variant caller, SVIM-asm [29]. This step is done for evaluation purposes because SVarp generates variant sequences instead of specific breakpoints. Then we compared generated variant breakpoints against the true call-set using BEDTools intersectBed [42] with 50% reciprocal overlap.

According to the results shown in Table 1, SVarp’s precision was excellent with values of 0.98 or larger across all categories. For shorter SVs, the sensitivity reaches values of 0.95 and above across SV categories. The most challenging SV classes are large insertions and duplications, where sensitivity is lower due to the more difficult alignment step as the SV length approaches the average read length of 10K. We believe this is a minor problem in real world scenarios because long SVs are much less common than shorter SVs within the scope of the read length.

**Table 1.** SVarp simulation results are given in the table. Precision and recall were calculated using the SVIM-asm predictions and are based on BEDtools 50% reciprocal overlap. We ran SVarp with a distance-threshold of 5000 and a minimum alignment score of 2000 (-d and -as parameters respectively).

|  |  | True SVs | Svtig | True | False | Miss | Precision | Recall |
| --- | --- | --- | --- | --- | --- | --- | --- | --- |
| **Deletion** | <1K | 276 | 305 | 268 | 0 | 8 | **1.00** | **0.97** |
|  | >1K | 276 | 327 | 262 | 6 | 14 | **0.98** | **0.95** |
| **Insertion** | <1K | 276 | 316 | 261 | 3 | 15 | **0.99** | **0.95** |
|  | >1K | 276 | 153 | 93 | 0 | 183 | **1.00** | **0.34** |
| **Duplication** | <1K | 276 | 303 | 264 | 3 | 12 | **0.99** | **0.96** |
|  | >1K | 276 | 261 | 214 | 1 | 62 | **1.00** | **0.78** |

We note that “Precision” in the table is based on SVIM-asm’s false calls compared to the true SV callset. It is difficult to attribute false predictions accurately in this scenario because each svtig outputted by SVarp may not correspond to one SV: It may either be a false SVarp prediction, a false SVIM-asm prediction, a SVIM-asm miss or multiple svtigs giving rise to a single SV (duplicate). So we cannot reliably conclude that the remaining svtigs are false calls of SVarp.

### 3.2 Long read ONT genome analysis

We used 10 ONT genomes (HG00268, HG00513, HG00731, HG02554, HG02953, NA12878, NA19129, NA19238, NA19331, NA19347) [43] to assess the performance of SVarp on real datasets. These have various depths of coverage ranging from 12-fold to 50-fold (average 20-fold), aligned to the HPRCv1.0 T2T-CHM13 pangenome graph. We selected these samples because they have accurate assemblies that we can use to verify our predictions [35].

First, using the widely analyzed NA12878 genome, we ran SVarp and aligned the generated svtigs against the assembly of NA12878 [35] to validate the generated svtigs (Figure 3). This is done using minimap2 [44], allowing up to 10% divergence (using parameter “-x asm10”) to align svtigs against the assembly. Then we calculate the sequence identity of each svtig if it has a single alignment. Otherwise, we use the sequence identity of the alignment with the largest mapping ratio, defined as the length of the aligned segment of the svtig divided by the total svtig length. A high quality svtig will both have high sequence similarity and a large mapping ratio, provided that the respective sequence context can be aligned. False svtigs will either not map to the assembly or have lower sequence identity. In this context, we did not observe any completely unmapped svtig. We also note that the assemblies used as ground truth have been generated using PacBio HiFi reads and are of high quality, but are not completely error-free. Overall, the svtigs show very good concordance with the assemblies, where 82% of the svtigs map to the assemblies with >85% identity and a mapping ratio of >85%. A fraction of 59% of all svtigs even meet more stringent criteria and map to assemblies with >95% identity and a mapping ratio >95%.

**Figure 3.**
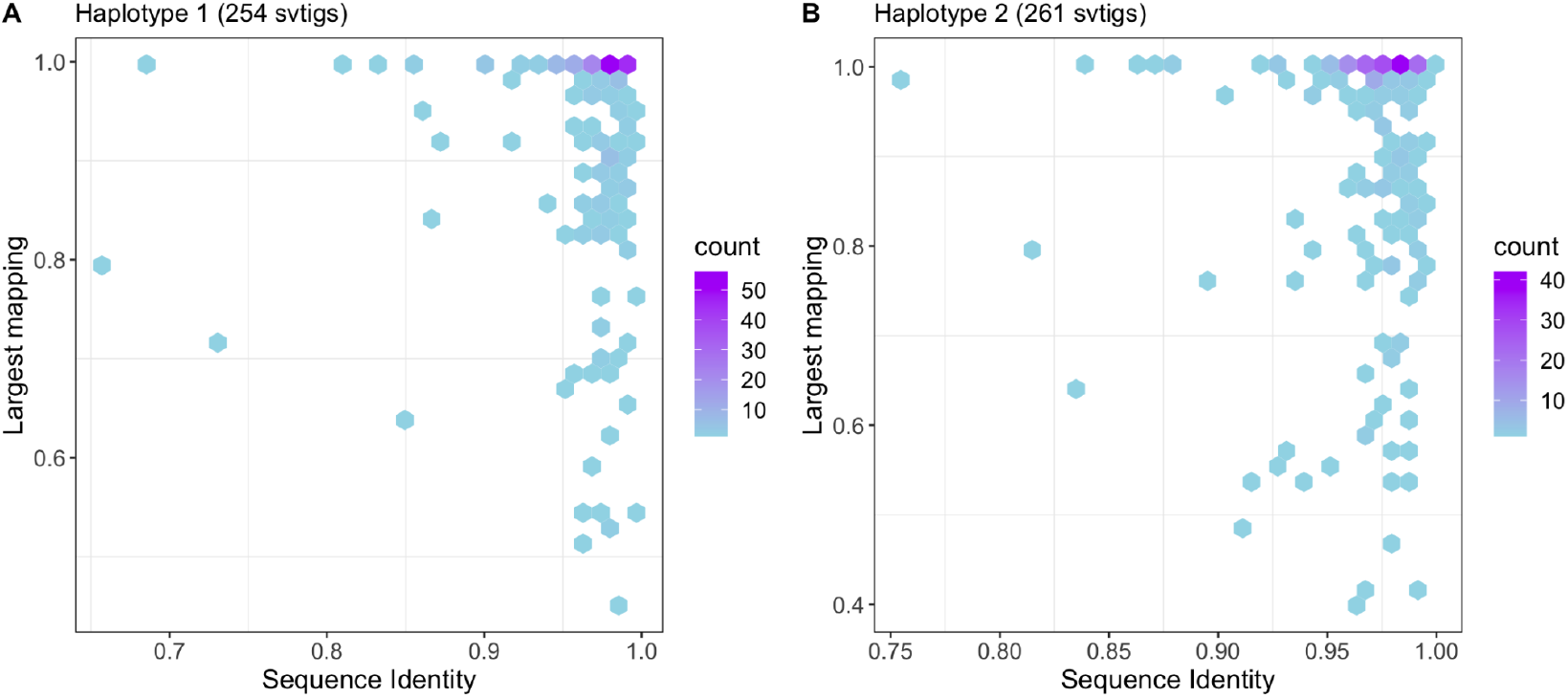
After running SVarp with NA12878 ONT genome, we aligned the output svtigs (254 for Haplotype 1 and 261 for Haplotype 2) against the HGSVC2 genome assembly for both haplotypes using minimap2 with parameter “-x asm10”. This plot shows the sequence identity for the largest mapping of each aligned svtig.

Then we ran SVarp on all ten ONT genomes and compared against the respective assemblies (Figure 4-A). Here, we define an svtig to be concordant if both its sequence identity of the largest alignment and the largest alignment itself are > 85%. For each sample, we divided the number of concordant svtigs by the total number of svtig predictions of SVarp and plotted the concordance ratio. We observe stable concordance values larger than 75% across all samples, while the number of svtigs shows a trend to grow with coverage. We attribute this to the increased chance of reaching the coverage cutoff with growing coverage.

**Figure 4.**
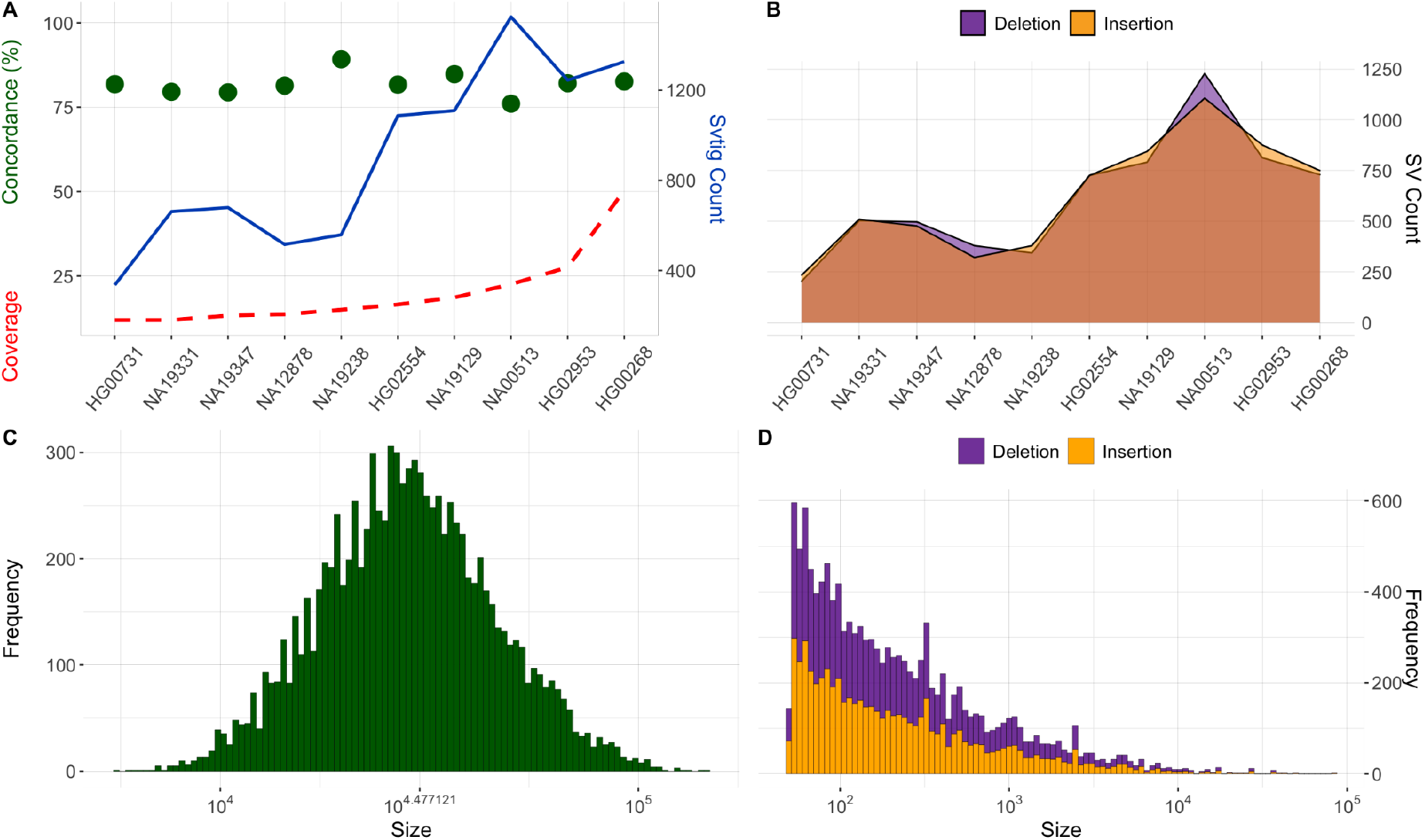
A) The relationship between concordance, coverage and svtig count are illustrated for the ONT samples. Concordance (green points) for each sample is calculated by aligning svtigs to the related assembly using minimap2 with parameter “-x asm10” and finding the sequence identity of the largest mapping for each svtig alignment. Then we calculate concordance ratio by summing the number of svtigs that have the largest mapping and sequence identity >85% and dividing by the SVarp’s svtig prediction count. This is done for both haplotypes separately and the plot shows the average. Blue line shows the total number of svtig predictions of each haplotype by SVarp and the red dotted line shows the coverage of each sample. B) Number of deletions (purple) and insertions (orange) generated by SVIM-asm relative to T2T reference genome, given the input svtigs generated by SVarp (using -d 100). C) Length distribution of svtigs generated by SVarp for the same genomes. D) Length distribution of deletions (purple) and insertions (orange) predicted by SVIM-asm.

Furthermore, we called SVs for each genome relative to the linear T2T reference, using SVIM-asm with SVarp’s svtig predictions as input. This produced 622 deletions and 623 insertions on average for the ten samples (Figure 4-B), yielding a similar trend to svtig counts shown in Figure 4-A. The average number of svtigs compared to the called SVs is slightly less, where an average of 904 svtigs per sample gives rise to on average 1245 SVs being called per sample, which mirrors what we observed forh simulations (also discussed in Section 3.1). We attribute this gap to multiple reasons. First, although we filtered duplicate calls of SVIM-asm, there are still multiple SVs within close proximity. We suspect that some of these are still duplicate predictions and SVIM-asm outputs them as distinct SVs. Second, we noticed that some large read clusters give rise to multiple SVs, where each of these might still be correct, which could occur in complex regions such as variable-number tandem repeats (VNTRx). In Figure 4-C and 4-D, we provide the size frequency distribution of svtigs and SVIM-asm generated SV breakpoints of insertions and deletions respectively. Of note, the number of insertions and deletions is balanced across the size spectrum, demonstrating a comparable ability to call both SV types for the shown length classes.

Finally, in order to further elucidate the effect of coverage on the accuracy and distance-threshold parameter, we used the highest coverage genome (HG00268) in our dataset and downsampled it to lower depths (Figure 5). For this experiment, we removed 70, 55, 40 and 25 percent of the reads in the fasta file randomly and aligned the remaining reads to the pangenome graph separately and generated the alignment GAF files. Then we called svtigs using SVarp. The distance threshold, depicted in Step 2 of Figure 1 with green and purple colors indicate that reads supporting adjacent SV breakpoints with proximity less than the threshold are merged (green ones) into the same cluster. Permitting a larger distance will have a tendency to create larger clusters and hence increases the chances that a cluster passes the read depth filter. On the one hand, larger values therefore increase the recall especially for lower coverage samples, but on the other hand, smaller values (such as -d 100) will give rise to fewer svtigs with higher reliability as shown in Figure 5.

**Figure 5.**
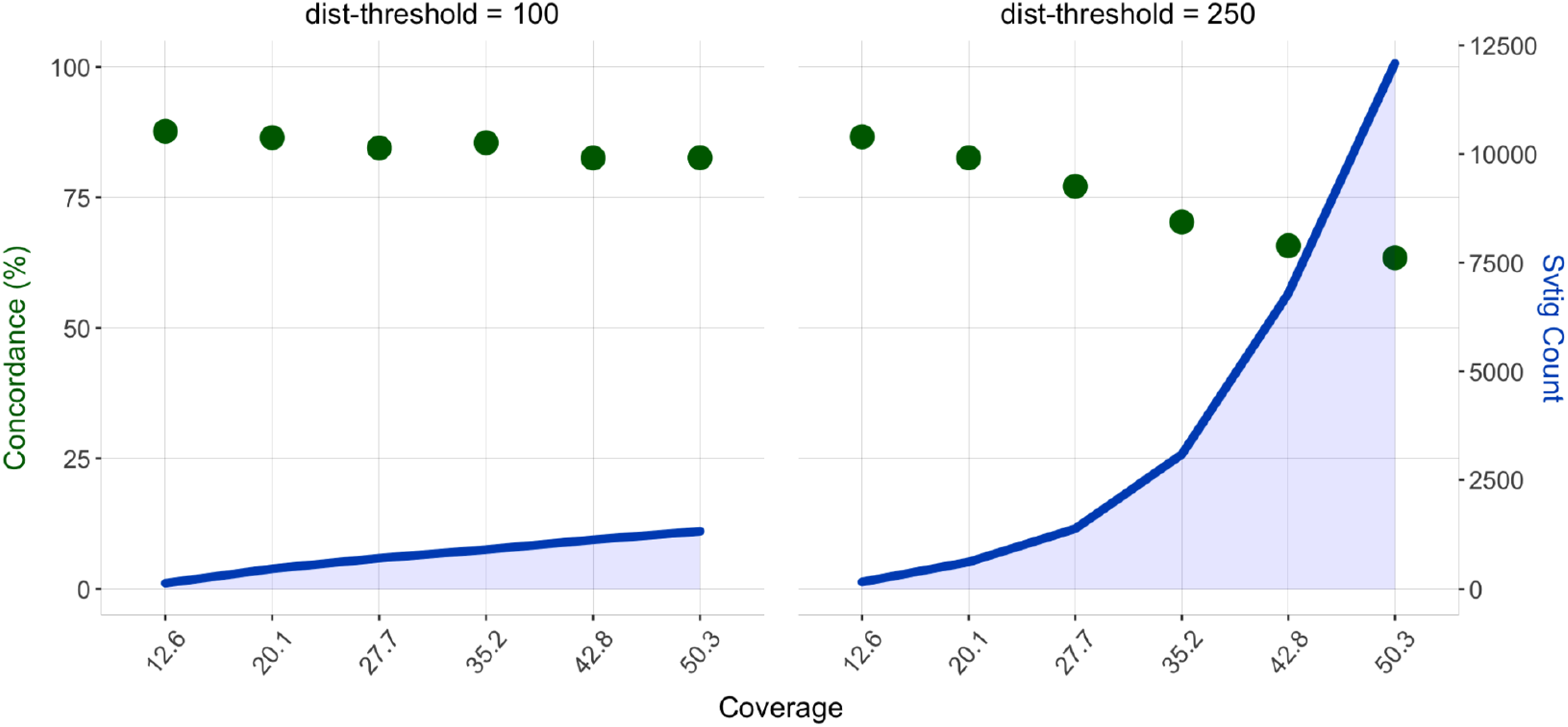
Downsampling experiment of the HG00268 genome, where we downsampled the original genome (50.3X relative to the pangenome graph) to lower depths to see the performance of SVarp in lower coverages. We use distance-threshold (-d) parameters of 100 (left) and 250 (right).

### 3.3 Runtime and memory

In order to evaluate the runtime and memory consumption of SVarp, we ran each of the downsampled HG00268 genomes described above using both distance-threshold of 100 and 250, in order to see the effect of increased svtig count (thus, higher coverage genomes) on the computational performance. This is done using the same computing resources for each genome with a single CPU core^1^. The x-axis in Figure 6 displays the increase in the number of svtigs, which is proportional to the number of assembly operations. Thus, the sharp increase with respect to the svtig count is due to increased runtime and memory in the local assembly process. Also, as the coverage increases, the number of reads that contribute to the cluster to be assembled increases, and so the runtime and memory. Overall, the computational resources are manageable and SVarp can even be run on a standard laptop for moderate coverage samples with standard distance-threshold value.

**Figure 6.**
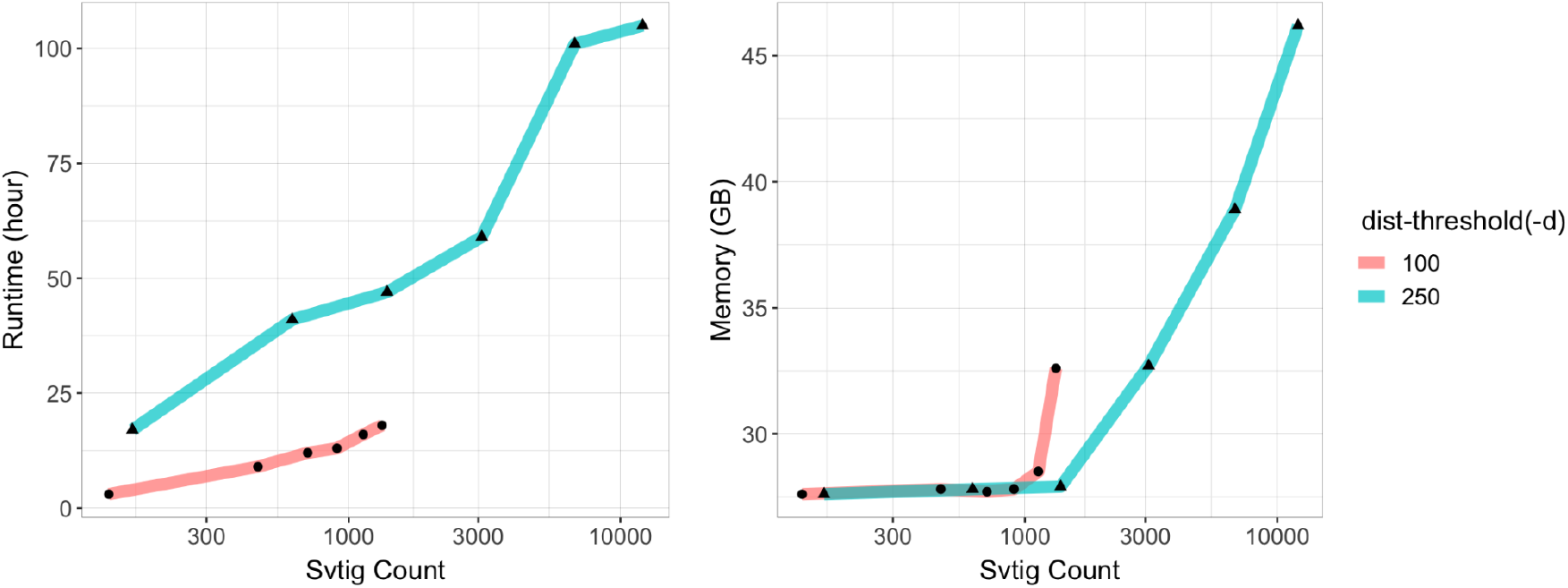
Runtime and memory consumption of the HG00268 genome downsampled to ∼43X, ∼35X, ∼28X, ∼20X, ∼13X from 50X shown with points on the line. Runtimes are given for a single CPU core. This is done for both distance-threshold of 100 and 250 with respect to the number of svtigs generated.

## 4. Conclusion and Discussion

The availability of the first human pangenome reference [4] entails the need for computational tools and algorithms to use these new resources. With SVarp, we close one important gap in the tool ecosystem and provide a method to discover SVs in a sample mapped to a pangenome graph. This will enable us to find additional SVs that currently escape detection, in particular SVs arising in alternative haplotype contexts present in the pangenome but not in a linear reference. Rather than outputting a VCF file of variant types and breakpoints, we generate phased variant sequences, which we call svtigs. This is a distinction from the classic linear reference based SV callers because we believe that svtigs will enable more flexible subsequent analysis. As they are not tied to one coordinate system, but instead bring enough sequence context to be mapped in whatever linear or pangenomic coordinate system is employed in a downstream application.

Results with simulations show that SVarp can detect SVs reliably (>=98% precision, >=95% sensitivity) with the exception of very large insertions that have size close to the average read length. These have less sensitivity due to alignment difficulty of corresponding reads. Even though large insertions are comparatively rare and increasing read lengths will mitigate this problem, we plan to address this limitation through an iterative read recruitment strategy in the future. We also tested our approach using real ONT samples and observed that generated svtigs have high concordance with the assemblies of the corresponding samples, that is, 82% of the svtigs map to the assemblies with >85% sequence identity and >85% mapping ratio. These include low coverage samples (50% of the samples have less than 15X depth of coverage) as well and the concordance is shown to be stable with variations in coverage.

As a future direction, we believe graph augmentation using the svtigs that SVarp outputs would be a promising approach to make the pangenome reference more complete. Such an augmented graph reference would enable re-genotyping in large population sequencing projects and could also serve as a workflow to include variants from disease cohorts not consented for public release. Until such workflows that operate completely on pangenomes have matured, the svtigs output by SVarp can be used to produce SV calls relative to a linear reference, as we demonstrated in our evaluation.

## Code and Data Availability

SVarp is implemented in C++ and is available under MIT license at https://github.com/asylvz/SVarp. HPRC T2T-CHM13 pangenome graph can be found at https://zenodo.org/record/6983934. GAF files of the ONT genomes can be found in https://ftp.1000genomes.ebi.ac.uk/vol1/ftp/data_collections/1KG_ONT_VIENNA/

## Acknowledgements

This work was supported, in part, by the MODS project funded from the programme “Profilbildung 2020” [grant no. PROFILNRW-2020–107-A], an initiative of the Ministry of Culture and Science of the State of North Rhine-Westphalia, by the German Federal Ministry of Education and Research (BMBF) (031L0184A), and by National Human Genome Research Institute of the National Institutes of Health under award number U24HG007497. The authors would like to thank Siegfried Schloissnig for making the ONT data set available. We acknowledge the computational support provided by the Centre for Information and Media Technology (ZIM) at the Heinrich Heine University Düsseldorf.

## Footnotes

1 Intel(R) Xeon(R) Gold 6136 CPU @ 3.00GHz with 48 CPUs and 196 GB RAM

## References

1. Lander ES, Linton LM, Birren B, Nusbaum C, Zody MC, Baldwin J, et al. Initial sequencing and analysis of the human genome. Nature. 2001;409:860–921.

2. Iqbal Z, Caccamo M, Turner I, Flicek P, McVean G. De novo assembly and genotyping of variants using colored de Bruijn graphs. Nat Genet. 2012;44:226–32.

3. Garrison E, Sirén J, Novak AM, Hickey G, Eizenga JM, Dawson ET, et al. Variation graph toolkit improves read mapping by representing genetic variation in the reference. Nat Biotechnol. 2018;36:875–9.

4. Liao W-W, Asri M, Ebler J, Doerr D, Haukness M, Hickey G, et al. A draft human pangenome reference. Nature. 2023;617:312–24.

5. Chin C-S, Behera S, Khalak A, Sedlazeck FJ, Sudmant PH, Wagner J, et al. Multiscale analysis of pangenomes enables improved representation of genomic diversity for repetitive and clinically relevant genes. Nat Methods [Internet]. 2023; Available from: 10.1038/s41592-023-01914-y

6. Ebert P, Audano PA, Zhu Q, Rodriguez-Martin B, Porubsky D, Bonder MJ, et al. Haplotype-resolved diverse human genomes and integrated analysis of structural variation. Science [Internet]. 2021;372. Available from: 10.1126/science.abf7117

7. Chaisson MJP, Sanders AD, Zhao X, Malhotra A, Porubsky D, Rausch T, et al. Multi-platform discovery of haplotype-resolved structural variation in human genomes. Nat Commun. 2019;10:1784.

8. Audano PA, Sulovari A, Graves-Lindsay TA, Cantsilieris S, Sorensen M, Welch AE, et al. Characterizing the Major Structural Variant Alleles of the Human Genome. Cell. 2019;176:663–75.e19.

9. Jun G, English AC, Metcalf GA, Yang J, Chaisson MJP, Pankratz N, et al. Structural variation across 138,134 samples in the TOPMed consortium [Internet]. bioRxiv. 2023 [cited 2023 Aug 31]. p. 2023.01.25.525428. Available from: 10.1101/2023.01.25.525428v1.abstract

10. Byrska-Bishop M, Evani US, Zhao X, Basile AO, Abel HJ, Regier AA, et al. High-coverage whole-genome sequencing of the expanded 1000 Genomes Project cohort including 602 trios. Cell. 2022;185:3426–40.e19.

11. Chiang C, Scott AJ, Davis JR, Tsang EK, Li X, Kim Y, et al. The impact of structural variation on human gene expression. Nat Genet. 2017;49:692–9.

12. Sudmant PH, Rausch T, Gardner EJ, Handsaker RE, Abyzov A, Huddleston J, et al. An integrated map of structural variation in 2,504 human genomes. Nature. 2015;526:75–81.

13. Alkan C, Coe BP, Eichler EE. Genome structural variation discovery and genotyping. Nat Rev Genet. 2011;12:363–76.

14. Wenger AM, Peluso P, Rowell WJ, Chang P-C, Hall RJ, Concepcion GT, et al. Accurate circular consensus long-read sequencing improves variant detection and assembly of a human genome. Nat Biotechnol. 2019;37:1155–62.

15. Vollger MR, Logsdon GA, Audano PA, Sulovari A, Porubsky D, Peluso P, et al. Improved assembly and variant detection of a haploid human genome using single-molecule, high-fidelity long reads. Ann Hum Genet. 2020;84:125–40.

16. Logsdon GA, Vollger MR, Eichler EE. Long-read human genome sequencing and its applications. Nat Rev Genet. 2020;21:597–614.

17. De Coster W, Weissensteiner MH, Sedlazeck FJ. Towards population-scale long-read sequencing. Nat Rev Genet. 2021;22:572–87.

18. Nurk S, Koren S, Rhie A, Rautiainen M, Bzikadze AV, Mikheenko A, et al. The complete sequence of a human genome. Science. 2022;376:44–53.

19. Li R, Li Y, Zheng H, Luo R, Zhu H, Li Q, et al. Building the sequence map of the human pan-genome. Nat Biotechnol. 2010;28:57–63.

20. Martiniano R, Garrison E, Jones ER, Manica A, Durbin R. Removing reference bias and improving indel calling in ancient DNA data analysis by mapping to a sequence variation graph. Genome Biol. 2020;21:250.

21. Brandt DYC, Aguiar VRC, Bitarello BD, Nunes K, Goudet J, Meyer D. Mapping Bias Overestimates Reference Allele Frequencies at the HLA Genes in the 1000 Genomes Project Phase I Data. G3. 2015;5:931–41.

22. Rausch T, Zichner T, Schlattl A, Stütz AM, Benes V, Korbel JO. DELLY: structural variant discovery by integrated paired-end and split-read analysis. Bioinformatics. 2012;28:i333–9.

23. Layer RM, Chiang C, Quinlan AR, Hall IM. LUMPY: a probabilistic framework for structural variant discovery. Genome Biol. 2014;15:R84.

24. Chen X, Schulz-Trieglaff O, Shaw R, Barnes B, Schlesinger F, Källberg M, et al. Manta: rapid detection of structural variants and indels for germline and cancer sequencing applications. Bioinformatics. 2016;32:1220–2.

25. Soylev A, Kockan C, Hormozdiari F, Alkan C. Toolkit for automated and rapid discovery of structural variants. Methods. 2017;129:3–7.

26. Sedlazeck FJ, Rescheneder P, Smolka M, Fang H, Nattestad M, von Haeseler A, et al. Accurate detection of complex structural variations using single-molecule sequencing. Nat Methods. 2018;15:461–8.

27. Jiang T, Liu Y, Jiang Y, Li J, Gao Y, Cui Z, et al. Long-read-based human genomic structural variation detection with cuteSV. Genome Biol. 2020;21:189.

28. Heller D, Vingron M. SVIM: structural variant identification using mapped long reads. Bioinformatics. 2019;35:2907–15.

29. Heller D, Vingron M. SVIM-asm: Structural variant detection from haploid and diploid genome assemblies. Bioinformatics. 2020;36:5519–21.

30. Li H, Feng X, Chu C. The design and construction of reference pangenome graphs with minigraph. Genome Biol. 2020;21:265.

31. Hickey G, Monlong J, Ebler J, Novak AM, Eizenga JM, Gao Y, et al. Pangenome graph construction from genome alignments with Minigraph-Cactus. Nat Biotechnol [Internet]. 2023; Available from: 10.1038/s41587-023-01793-w

32. Rautiainen M, Marschall T. GraphAligner: rapid and versatile sequence-to-graph alignment. Genome Biol. 2020;21:253.

33. Rakocevic G, Semenyuk V, Lee W-P, Spencer J, Browning J, Johnson IJ, et al. Fast and accurate genomic analyses using genome graphs. Nat Genet. 2019;51:354–62.

34. Sirén J, Monlong J, Chang X, Novak AM, Eizenga JM, Markello C, et al. Pangenomics enables genotyping of known structural variants in 5202 diverse genomes. Science. 2021;374:abg8871.

35. Ebler J, Ebert P, Clarke WE, Rausch T, Audano PA, Houwaart T, et al. Pangenome-based genome inference allows efficient and accurate genotyping across a wide spectrum of variant classes. Nat Genet. 2022;54:518–25.

36. Martin M, Patterson M, Garg S, Fischer SO, Pisanti N, Klau GW, et al. WhatsHap: fast and accurate read-based phasing [Internet]. bioRxiv. 2016 [cited 2023 Sep 8]. p. 085050. Available from: 10.1101/085050v2.full

37. Ruan J, Li H. Fast and accurate long-read assembly with wtdbg2. Nat Methods. 2020;17:155–8.

38. Marco-Sola S, Eizenga JM, Guarracino A, Paten B, Garrison E, Moreto M. Optimal gap-affine alignment in O(s) space. Bioinformatics [Internet]. 2023;39. Available from: 10.1093/bioinformatics/btad074

39. Marco-Sola S, Moure JC, Moreto M, Espinosa A. Fast gap-affine pairwise alignment using the wavefront algorithm. Bioinformatics. 2021;37:456–63.

40. Bolognini D, Sanders A, Korbel JO, Magi A, Benes V, Rausch T. VISOR: a versatile haplotype-aware structural variant simulator for short-and long-read sequencing. Bioinformatics. 2020;36:1267–9.

41. Li H. Minigraph pangenome graphs for HPRC year-1 samples [Internet]. Zenodo; 2022. Available from: 10.5281/zenodo.6499594

42. Quinlan AR, Hall IM. BEDTools: a flexible suite of utilities for comparing genomic features. Bioinformatics. 2010;26:841–2.

43. Asparuhova, M., Ebler, J., Hüther, P., Korbel, J., Marschall, T., Pani, S., Rausch, T., Rodríguez-Martín, B., Schloissnig, S., Söylev, A., & Tsapalou, V. SV analysis of the 1019 samples of the 1KG-ONT panel (v1.0.0) [Internet]. 2023. Available from: https://zenodo.org/records/10418434

44. Li H. Minimap2: pairwise alignment for nucleotide sequences. Bioinformatics. 2018;34:3094–100.

