## Appendix for "SVarp: pangenome-based structural variant discovery"

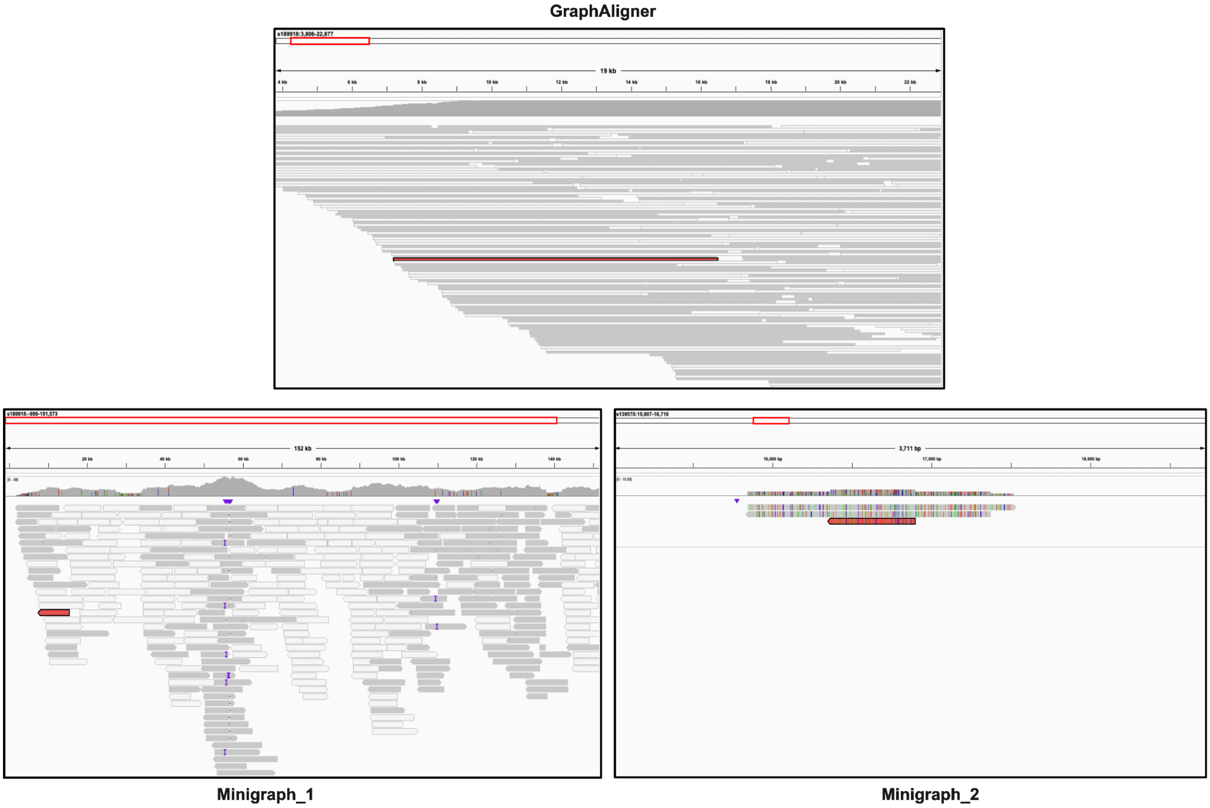


**Figure S1.** Alignment of a sample read (C1_H2_10586, shown in red) to HPRCv1.0 T2T-CHM13 Minigraph pangenome graph [1,2] using GraphAligner [3] (top) and Minigraph [4] (bottom). The read of length 9303 bps has a single mapping with GraphAligner, however Minigraph chops it into two alignments (Minigraph_1 and Minigraph_2). We used Lorax [5] to convert pangenome alignments into BAM format [6] to visualize using IGV [7].

**Table S1.** Comparison of GraphAligner and Minigraph using NA12878 ONT dataset (~14X) [8] and HPRCv1.0 T2T-CHM13 pangenome reference (using 24 cores and 24 threads)

| Aligner | Time (h:m) | Memory (GB) |
|---|---|---|
| GraphAligner | 71:27 | 28 |
| Minigraph | 1:57 | 26 |

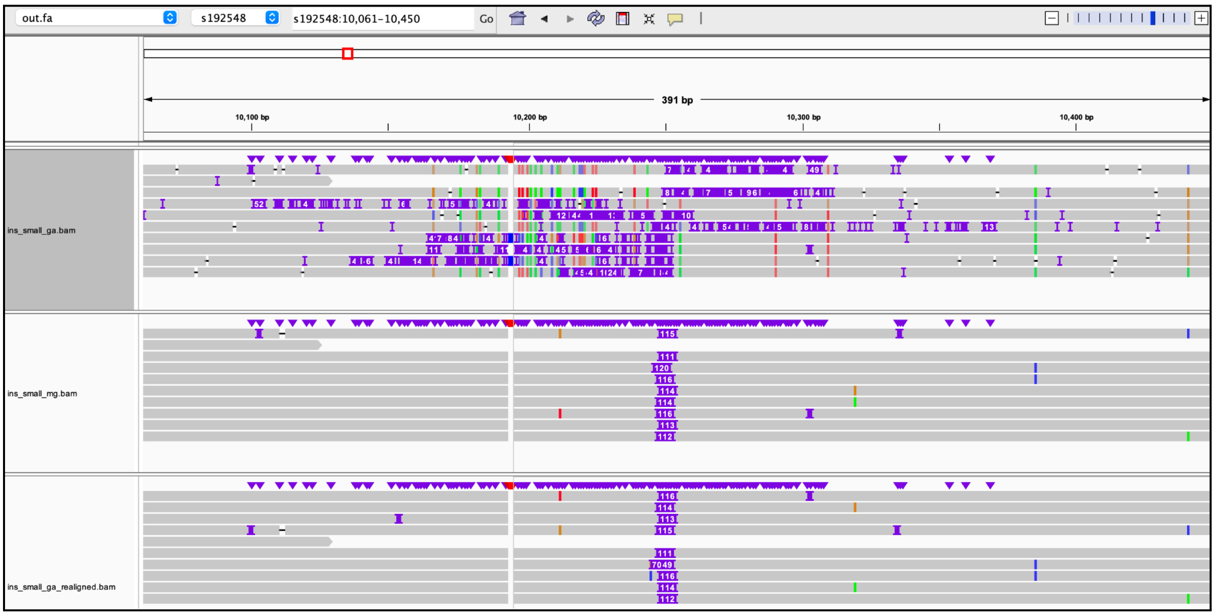


**Figure S2.** Alignment of a sample read to HPRCv1.0 T2T-CHM13 Minigraph pangenome graph using GraphAligner (top) and Minigraph (middle). We used Lorax to convert pangenome alignments into BAM alignments to visualize using IGV. Although Minigraph finds the insertion accurately, GraphAligner fails to show the correct position of the variant within the alignment. However, by using the WFA [9] algorithm and realigning the GraphAligner alignments, the problem is resolved as shown (bottom).

### References

1. Li H. Minigraph pangenome graphs for HPRC year-1 samples [Internet]. Zenodo; 2022. Available from: https://doi.org/10.5281/zenodo.6499594

2. Liao W-W, Asri M, Ebler J, Doerr D, Haukness M, Hickey G, et al. A draft human pangenome reference. Nature. 2023;617:312–24.

3. Rautiainen M, Marschall T. GraphAligner: rapid and versatile sequence-to-graph alignment. Genome Biol. 2020;21:253.

4. Li H, Feng X, Chu C. The design and construction of reference pangenome graphs with minigraph. Genome Biol. 2020;21:265.

5. Rausch T, Snajder R, Leger A, Simovic M, Giurgiu M, Villacorta L, et al. Long-read sequencing of diagnosis and post-therapy medulloblastoma reveals complex rearrangement patterns and epigenetic signatures. Cell Genom. 2023;3:100281.

6. Li H, Handsaker B, Wysoker A, Fennell T, Ruan J, Homer N, et al. The Sequence Alignment/Map format and SAMtools. Bioinformatics. 2009;25:2078–9.

7. Robinson JT, Thorvaldsdóttir H, Winckler W, Guttman M, Lander ES, Getz G, et al. Integrative genomics viewer. Nat Biotechnol. 2011;29:24–6.

8. Asparuhova, M., Ebler, J., Hüther, P., Korbel, J., Marschall, T., Pani, S., Rausch, T., Rodríguez-Martín, B., Schloissnig, S., Söylev, A., & Tsapalou, V. SV analysis of the 1019 samples of the 1KG-ONT panel (v1.0.0) [Internet]. 2023. Available from: https://zenodo.org/records/10418434

9. Marco-Sola S, Moure JC, Moreto M, Espinosa A. Fast gap-affine pairwise alignment using the wavefront algorithm. Bioinformatics. 2021;37:456–63.
